# A conformational checkpoint between DNA binding and cleavage by CRISPR-Cas9

**DOI:** 10.1101/122242

**Authors:** Yavuz S. Dagdas, Janice S. Chen, Samuel H. Sternberg, Jennifer A. Doudna, Ahmet Yildiz

**Affiliations:** Biophysics Graduate Group, University of California, Berkeley, California 94720, USA; Department of Molecular and Cell Biology, University of California, Berkeley, California, 94720, USA; Department of Chemistry, University of California, Berkeley, California 94720, USA; Howard Hughes Medical Institute, University of California, Berkeley, California 94720, USA; Physical Biosciences Division, Lawrence Berkeley National Laboratory, Berkeley, California 94720, USA; Department of Physics, University of California, Berkeley, California 94720, USA

## Abstract

The Cas9 endonuclease is widely utilized for genome engineering applications by programming its single-guide RNA and ongoing work is aimed at improving the accuracy and efficiency of DNA targeting. DNA cleavage of Cas9 is controlled by the conformational state of the HNH nuclease domain, but the mechanism that governs HNH activation at on-target DNA while reducing cleavage activity at off-target sites remains poorly understood. Using single-molecule FRET, we identified an intermediate state of S. pyogenes Cas9, representing a conformational checkpoint between DNA binding and cleavage. Upon DNA binding, the HNH domain transitions between multiple conformations before docking into its active state. HNH docking requires divalent cations, but not strand scission, and this docked conformation persists following DNA cleavage. Sequence mismatches between the DNA target and guide RNA prevent transitions from the checkpoint intermediate to the active conformation, providing selective avoidance of DNA cleavage at stably bound off-target sites.

---

The RNA-guided endonuclease Cas9 is responsible for recognizing, unwinding and cutting double-stranded DNA targets as part of the type II CRISPR-Cas (Clustered Regularly Interspaced Short Palindromic Repeats–CRISPR-associated) adaptive immune system (*1-4*). DNA target recognition requires a short PAM sequence (5’-NGG-3’ for *Streptococcus pyogenes* Cas9) (*5*) and complementary base pairing with the 20 nt targeting sequence of the guide RNA. Cleavage of the target (TS) and non-target (NTS) strands is mediated by the conserved HNH and RuvC nuclease domains, respectively (*6, 7*). By manipulating the sequence of the single-guide RNA (sgRNA), Cas9-sgRNA can be programmed to target any DNA sequence flanked by a PAM (*6, 8-10*), making it a powerful genome editing tool. However, promiscuous cleavage of off-target sites remains a major challenge (*5, 11-14*). Efforts are in progress (*15, 16*) to enhance the specificity of DNA targeting by Cas9 for potential therapeutic applications (*17, 18*), which requires detailed mechanistic understanding of substrate-dependent activation of Cas9 for DNA cleavage.

A previous bulk FRET study revealed that the HNH conformation regulates DNA cleavage activity of both nuclease domains and is highly sensitive to RNA-DNA complementarity (*12*). After binding to a DNA target, *S. pyogenes* Cas9 undergoes large conformational rearrangements that enable the HNH active site to hydrolyze the TS scissile phosphate (*19*) (Fig. 1A). However, none of the published structures of Cas9 have captured the HNH domain at the cleavage site (*19-22*) and the molecular cues that govern transitions to the active conformation are not well understood. Because bulk methods are insensitive to underlying dynamics, it stands unresolved whether the HNH domain only transiently switches to the active conformation or stably docks onto the active site for DNA cleavage. Furthermore, Cas9 cleaves only a subset of off-target sites to which it binds (*23, 24*), but it remains unclear whether HNH conformational dynamics play a direct role in decoupling between off-target binding and cleavage.

**Fig. 1.**
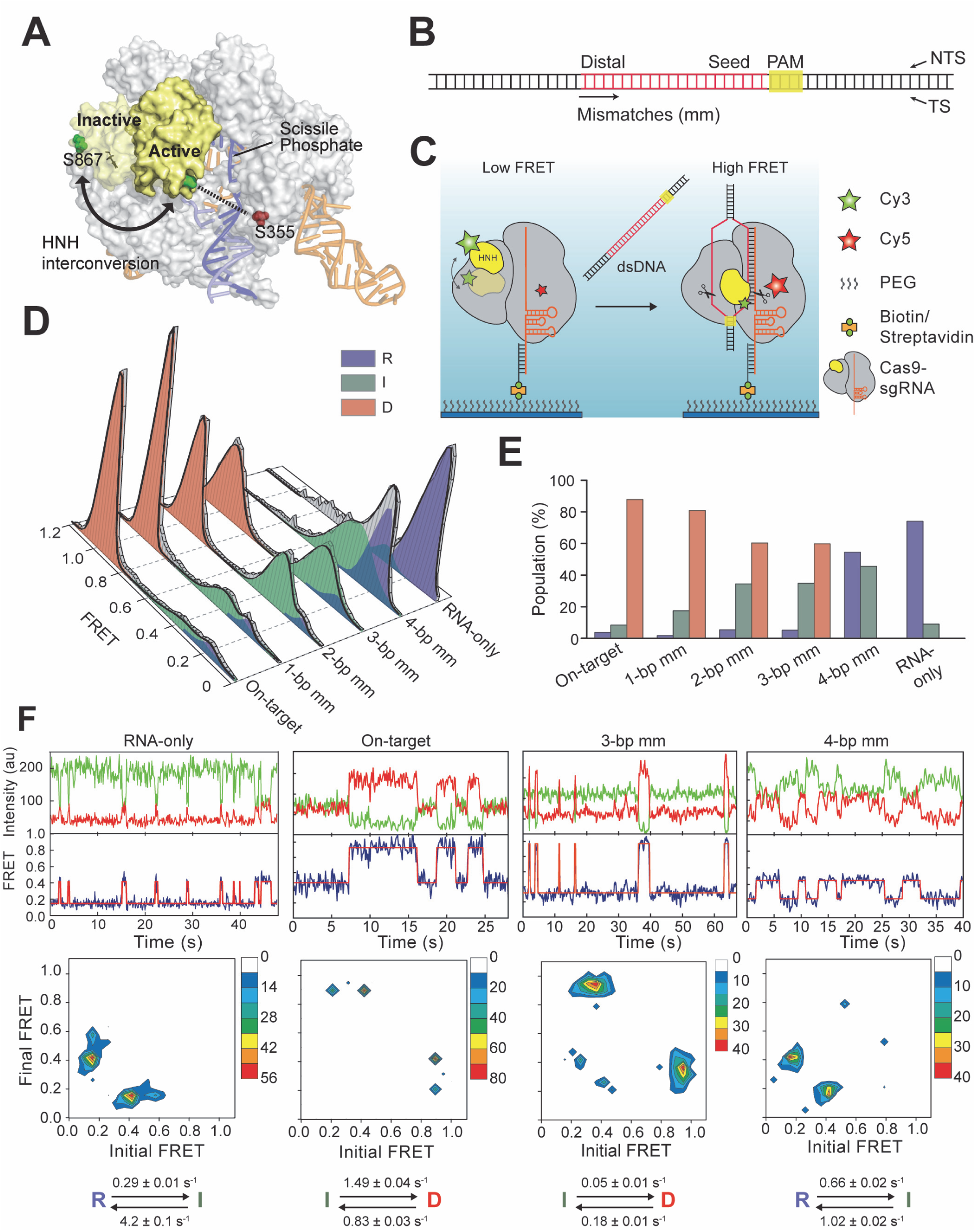
HNH conformational dynamics reveal a distinct I state as a function of PAM-distal mismatches. (**A**) Model shows HNH labeling sites under different conformations of Cas9, using sgRNA-bound (4ZT0) and dsDNA-bound (5F9R) structures. The cysteine-light Cas9 construct is labeled with Cy3 and Cy5 at S867C and S355C positions. (**B**) Cas9 was incubated with 55-bp long dsDNA substrates that include PAM and target sequences. Mismatches were introduced at the PAM distal site. (**C**) DNA binding to Cas9 results in HNH interconversion, determined by a transition from a low to high FRET state. Scissors show the DNA cleavage sites. (**D**) Steady-state smFRET histograms in the absence and presence of 200 nM DNA targets. A multi-Gaussian fitting (black curve) reveals D, I and R states of HNH. (**E**) The percentage of distinct conformational states of Cas9 for various DNA substrates. (**F**) Representative time traces (top), transition density plots (TDPs, middle) and rates of the major transitions in TDPs (bottom) for various DNA substrates.

To address these questions, we monitored the conformational rearrangements of the HNH domain during DNA binding and cleavage using single-molecule fluorescence resonance energy transfer (smFRET). A cysteine-light *S. pyogenes* Cas9 variant was labeled at positions S355C and S867C with Cy3 and Cy5 dyes (*12*) (Fig. 1A), which remained fully functional for DNA cleavage (*12*) (fig. S1). Cas9 was pre-assembled with the sgRNA and DNA substrate (Fig. 1B and table S1), surface-immobilized via the sgRNA, and imaged under total internal reflection (TIR) excitation (Fig. 1C). Steady-state smFRET measurements revealed distinct conformations of Cas9 (fig. S2). Without the DNA substrate, the majority of the complexes exhibited a FRET efficiency (E_FRET_) of 0.19 ± 0.02 (mean ± SEM), referred to as the RNA-bound (R) state (Fig. 1D). Addition of 200 nM on-target DNA resulted in a near-complete loss of the R population and the appearance of a new population at 0.97 ± 0.01 E_FRET_, referred to as the docked (D) state (Fig. 1D). The D state was not observed when Cas9-sgRNA was mixed with non-target or no-PAM DNA (fig. S1), suggesting that this conformation requires stable RNA-DNA base pairing (*25*).

We next assembled Cas9-sgRNA with off-target DNA substrates containing 1-, 2-, 3-, or 4-bp mismatches (mm) at the PAM-distal end of the target (Fig. 1B) to understand how sensing of the RNA/DNA heteroduplex affects the HNH conformation (*19*). When Cas9 was premixed with these substrates, a third peak emerged at 0.34 ± 0.03 E_FRET_ (Fig. 1D), referred to as the intermediate (I) state. As the number of mismatches was increased, we observed a steady decrease in the D population, coupled with an increase in I and R populations. Remarkably, Cas9 was entirely unable to attain its D conformation with a 4-bp mm DNA (Fig. 1, D and E), consistent with the inability of Cas9 to cleave this substrate (*12*). We concluded that the D conformation represents the active docked state for DNA cleavage. E_FRET_ of the R, I and D states are consistent with the distance between the labeled positions from available structures and the predicted cleavage-competent conformation of Cas9 (*26, 27*) (table S2). Similar results were obtained with a reciprocal Cas9 variant (*12*) (fig. S3), verifying that our conclusions are not markedly affected by the dye labeling positions.

We analyzed individual smFRET trajectories to address whether HNH activation is completely prohibited or the residence time in the active conformation is reduced at off-target sites. In the absence of DNA, ∼50% of Cas9-sgRNA complexes displayed a stable R conformation (fig. S4), while the remaining 50% briefly visits the I state (Fig. 1F and fig. S5), indicating that this conformation is not stable without DNA. When Cas9 was bound to its on-target (fig. S6), the majority (90%) of the complexes maintained a stable D conformation (fig. S4) and only 3% displayed dynamic transitions between D and I states (Fig. 1F). In comparison to an on-target, a larger percentage (35%) of complexes underwent transitions between I and D states at a ∼100-fold slower rate with a 3-bp mm DNA. The complexes bound to a 4-bp mm DNA transitioned only between R and I states, whereas transitions to the D state were not observed (Fig. 1F). These results suggest that the I state serves as the conformational checkpoint of RNA-DNA complementarity before the HNH domain transitions to the D state.

To test the checkpoint hypothesis, we determined the real-time kinetics of HNH domain activation immediately upon binding to surface-immobilized DNA (Fig. 2A). The complexes were in the R state when initially bound to DNA (Fig. 2B). Individual trajectories recorded at 100 Hz revealed that Cas9 visits the I state (τ = 34 ms) between initial landing and transitioning to the D state (Fig. 2B), consistent with this state serving as the checkpoint between DNA binding and cleavage. The first transition to the D state occurs rapidly (0.7 s^−1^) after initial binding to an on-target DNA (Fig. 2, C, D and G). 95% of the complexes stably remained in a D conformation (Fig. 2C and fig. S7), and reversible transitions to the I and R states were rarely observed. With a 3-bp mm DNA, the first transition to D occurs at a ∼10-fold slower rate relative to an on-target DNA (Fig. 2, E, F and G) and the majority of complexes underwent reversible transitions after reaching the D state (Fig. 2D and fig. S7).

**Fig. 2.**
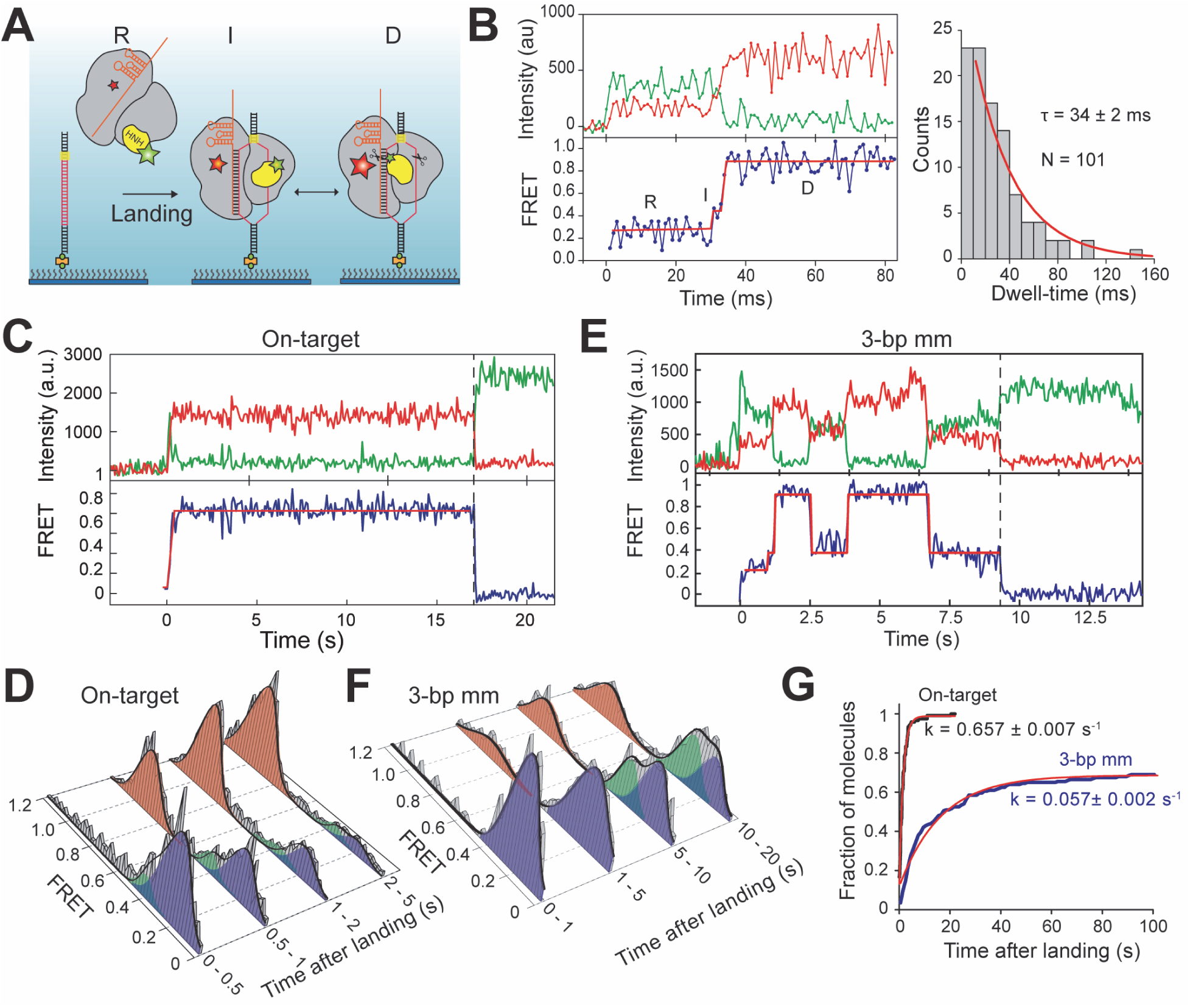
The real-time kinetics of HNH domain activation immediately after DNA binding. (**A**) Schematic for observation of Cas9 conformational dynamics upon landing onto surface-immobilized DNA. (**B**) (Left) A representative smFRET trajectory recorded at 100 Hz shows a brief visit to the I state between initial binding to an on-target DNA and transitioning into the D state. (Right) Single exponential fit (red curve) to the dwell time distribution of the I state reveals its lifetime (τ, ±95% confidence interval). (**C and E**) A representative smFRET trajectory at 10Hz of Cas9 after landing to an on-target and 3-bp mm DNA. t = 0 s and dotted vertical lines represent time of landing and acceptor photobleaching, respectively. (**D and F**) Time-dependent changes in the conformational distribution of Cas9 after landing onto an on-target and 3-bp mm DNA. (**G**) Cumulative distribution of first transition to the D conformation after landing onto a DNA. Red curves show fit to a single exponential function (±95% confidence interval).

Next, we tested the roles of divalent cations and nuclease activity on conformational activation of the HNH domain. DNA cleavage activity of Cas9 requires Mg^2+^ at the catalytic site (*6*), yet it remains stably bound to the DNA target without Mg^2+^ (fig. S8). It was unclear whether Mg^2+^ enables transition to the D state or stabilizes the D state via interactions with the active site and DNA phosphate backbone (*19*). We observed that Cas9 is completely unable to transition to the D state without divalent cations (Fig. 3A, fig. S1). At low (10 μM) Mg^2^+, 29% of molecules were in the D conformation (Fig. 3A), and 50% of trajectories revealed I to D transitions (fig. S9). At 1-5 mM Mg^2^+, Ca^2^+ or Co^2^+, >85% of complexes populated the D state, demonstrating that divalent cations lower the threshold energy for the HNH domain to move to its active state. We observed the same divalent cation dependent docking for catalytically-dead dCas9 (Fig. 3B). Furthermore, Co^2+^ is unable to support DNA cleavage (fig. S8), but enabled transitions to the D state (Fig. 3A) (*6*), demonstrating that docking of the HNH domain is independent of nuclease activity.

**Fig. 3.**
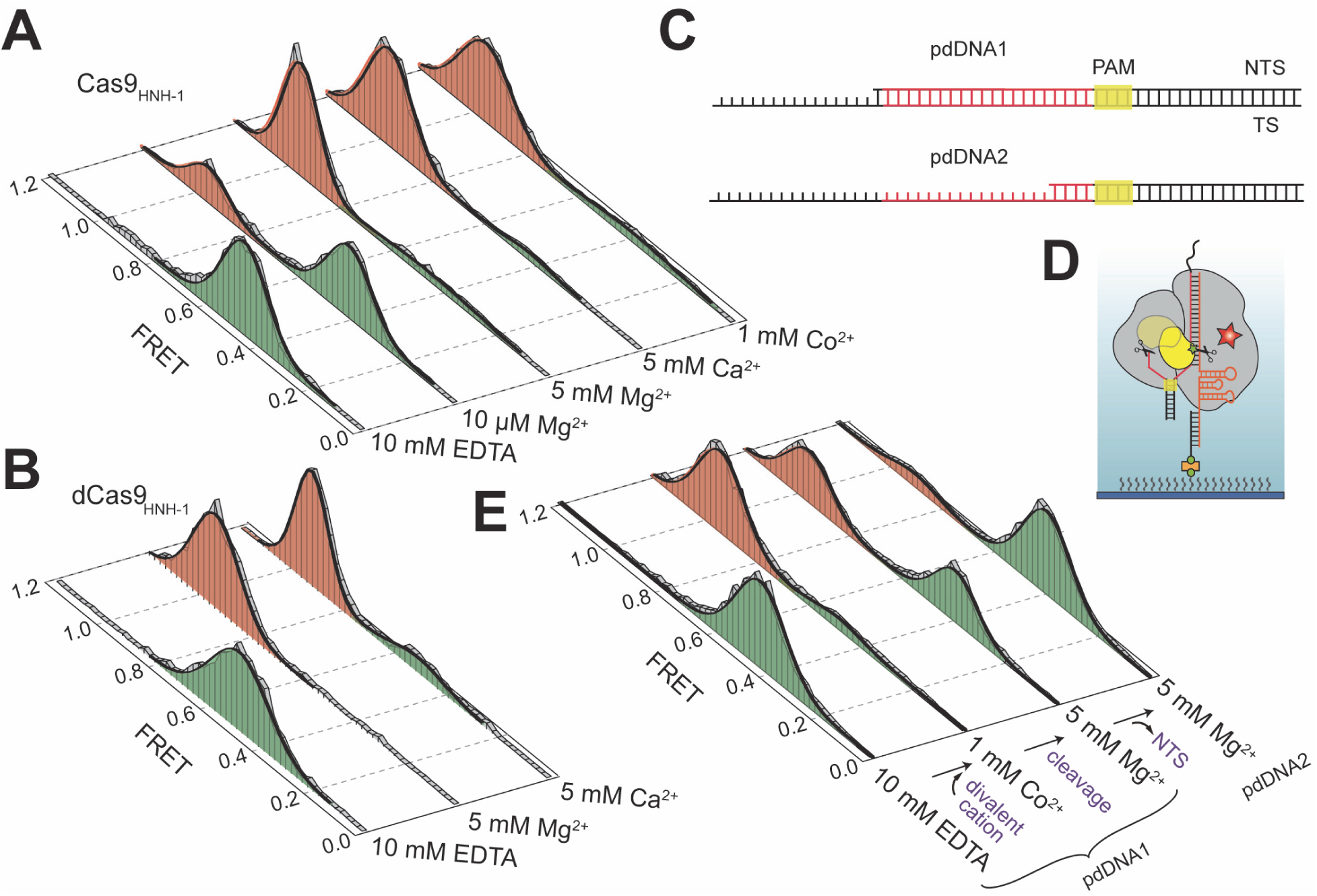
Docking of HNH to its active state requires divalent cation. (**A and B**) smFRET histograms of Cas9 and dCas9 bound to on-target dsDNA in the absence and presence of a divalent cation. (**C**) The on-target DNA was truncated at the 5’-end of the NTS one base after the target sequence (pdDNA1) and 4 bases after PAM (pdDNA2). (**D**) pdDNA1 and 2 mimic the DNA substrates after cleavage and NTS release, respectively. (**E**) smFRET histograms of Cas9 bound to pdDNAs in the absence and presence of a divalent cation.

To explore the stability of the D conformation after NTS release, we truncated to the 5’-end of the spacer in the NTS (pdDNA1) to prevent base pairing beyond the target region and enable dissociation of the NTS 5’-end from the complex after cleavage (Fig. 3, C and D) (*28*). The complexes bound to pdDNA1 transitioned to the D state upon addition of Co^2^+. Unlike an on-target dsDNA, in which the NTS remains bound to the complex post-cleavage, initiating the cleavage of pdDNA1 by Mg^2+^ addition reduced the stability of the D conformation (Fig. 3E). Furthermore, when Cas9 was bound to a distinct substrate that lacks the entire 5’-end of NTS upstream of the cleavage site (pdDNA2), the majority of complexes transitioned to the I conformation (Fig. 3, C to E). Because the HNH domain reverts back to the I state upon NTS release and cannot dock in the absence of the NTS, the 5’end of the NTS is necessary for stabilizing the D state (*29*).

Finally, we designed a single-molecule stopped flow assay to determine whether stable docking of HNH is rate limiting for the DNA cleavage activity (Fig. 4A). In this assay, pdDNA1 was labeled with Cy5 at the 5’-end of NTS, preassembled with Cas9 in 10 mM EDTA, and then complexes were immobilized via the sgRNA (Fig. 4B). Replacement of EDTA with 5 mM Mg^2+^ resulted in disappearance of 80% of the Cy5 spots at a rate of 0.16 s^−1^ (Fig. 4C and movie S1), which represents DNA cleavage and NTS release. Because docking of the HNH domain occurs ∼4-fold faster than DNA cleavage, it is not rate-limiting for cleavage of the on-target DNA. However, when Cas9 was preassembled with a 3-bp mm DNA, HNH docking and DNA cleavage (0.038 s^−1^, Fig. 4C and movie S2) occurred at comparable rates, suggesting that HNH docking is a rate-limiting step for the cleavage of this substrate.

**Fig. 4.**
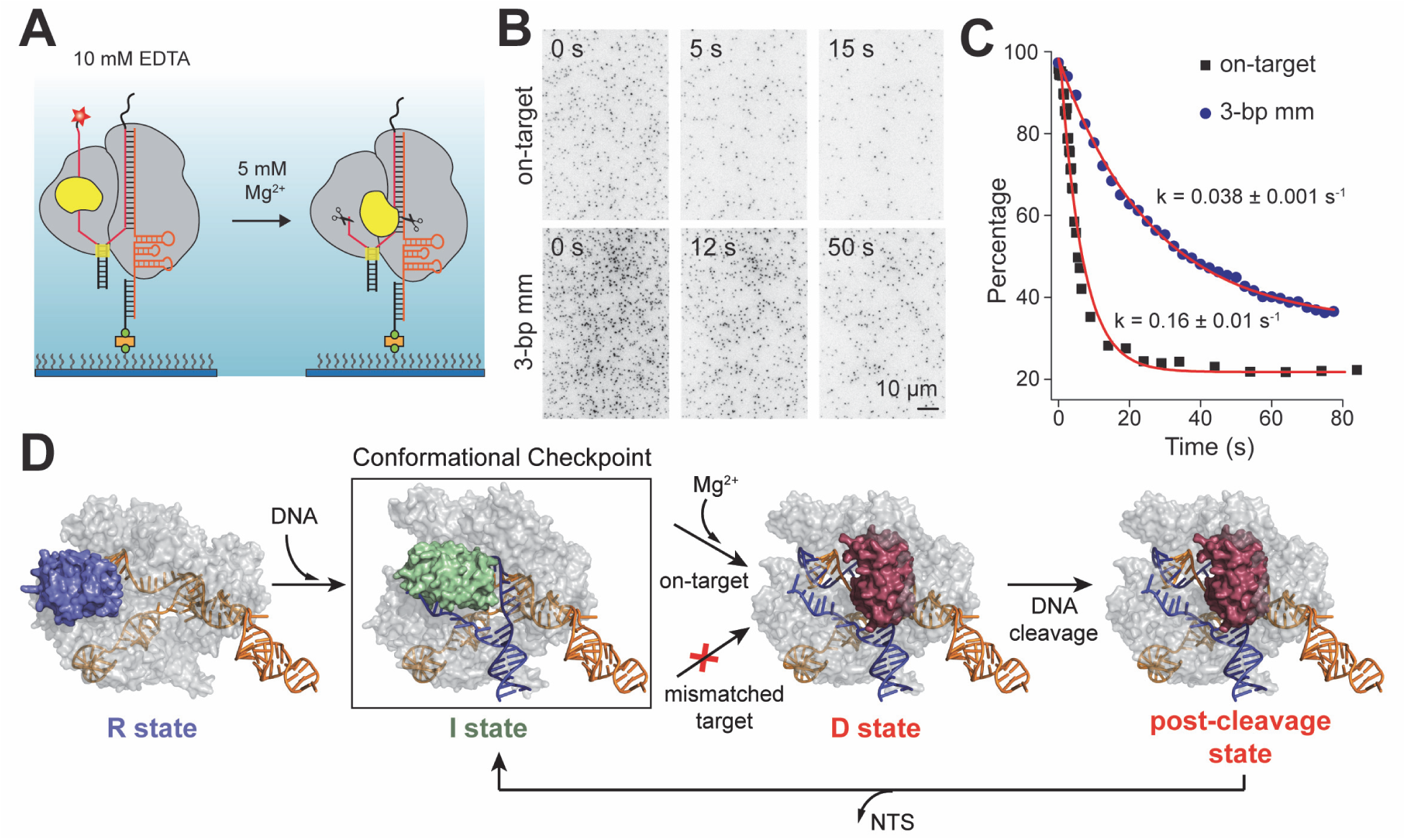
HNH docking is rate limiting for the cleavage of off-target, but not the on-target DNA. (**A**) The DNA cleavage activity of Cas9 was initiated by replacing EDTA in assay solution with 5 mM Mg^2+^ and monitored by dissociation of the Cy5-labeled NTS upon cleavage. (**B**) Still images of Cy5-labeled DNA bound to surface-immobilized Cas9 after Mg^2+^ addition (t = 0 s). (**C**) The percentage of Cy5 spots remain at the surface after Mg^2+^ flow. Red curves represent fit to single exponential decay (mean ± 95% confidence interval). (**D**) Model for substrate-dependent HNH activation. Without DNA, the HNH nuclease occupies the R state (blue). DNA binding triggers transition to the catalytically-inactive I state (green), which serves as a conformational checkpoint between DNA binding and cleavage. In the presence of divalent cations, recognition of an on-target locks the HNH nuclease in the catalytically-active D state, and Cas9 remains in this conformation post-cleavage. The D state is destabilized after NTS release. Transitions to the D state are prohibited when Cas9 is bound to certain off-target DNA substrates (red cross).

In this study, we detected conformational activation of Cas9 for DNA cleavage (Fig. 4D). Upon binding to an on-target DNA, the HNH domain transitions from the RNA-bound to catalytically-active D conformation. HNH remains docked to the catalytic site post-cleavage (*5, 13, 14, 28*) and this conformation is destabilized upon NTS release. Binding to a divalent cation is a prerequisite for docking into the HNH active state. These results explain why the active conformation of the HNH domain was not observed without the full dsDNA substrate or in the absence of divalent cations (*19-22*). We speculate that it may be captured with wild-type Cas9 in Co^2+^ or with dCas9 in Mg^2^+.

In the presence of mismatched targets, the emergence of the catalytically-inactive I state highlights a conformational checkpoint through which the HNH nuclease must pass to occupy the D conformation and achieve DNA cleavage (Fig. 4D). The I conformation also explains how cleavage activity is reduced when Cas9 binds to off-target sites (*12*). PAM-distal mismatches favor transitions into the I state and docking into the active state is blocked when complementarity drops below a threshold. Together, these results demonstrate the inherent conformational specificity of Cas9, and provide a structural and kinetic explanation for the decoupling of DNA binding and cleavage. The energetic barrier regulating access to the D state may be exploited with targeted mutagenesis, and the smFRET approach developed here can be a powerful strategy to characterize the intrinsic fidelity of Cas9 for genome editing applications.

## Acknowledgments

We would like to thank V. Belyy and B. LaFrance for helpful discussions and revision of our manuscript, and T. Ha for the data acquisition software. JSC is supported by the National Science Foundation Graduate Research Fellowship. This work has been supported by NIH (GM094522 (AY), NSF (MCB-1055017 (AY) and MCB-1244557 (JAD)) and Howard Hughes Medical Institute (JAD).

